# Identifying size-dependent toxin sorting in bacterial outer membrane vesicles

**DOI:** 10.1101/2023.05.03.539273

**Authors:** Aarshi N. Singh, Justin B Nice, Angela C. Brown, Nathan J. Wittenberg

## Abstract

Gram-negative bacteria produce outer membrane vesicles (OMVs) that play a critical role in cell-cell communication and virulence. Despite being isolated from a single population of bacteria, OMVs can exhibit heterogeneous size and toxin content, which can be obscured by assays that measure ensemble properties. To address this issue, we utilize fluorescence imaging of individual OMVs to reveal size-dependent toxin sorting. Our results showed that the oral bacterium *Aggregatibacter actinomycetemcomitans (A*.*a*.) produces OMVs with a bimodal size distribution, where larger OMVs were much more likely to possess leukotoxin (LtxA). Among the smallest OMVs (< 100 nm diameter), the fraction that are toxin positive ranges from 0-30%, while the largest OMVs (> 200 nm diameter) are between 70-100% toxin positive. Our single OMV imaging method provides a non-invasive way to observe OMV surface heterogeneity at the nanoscale level and determine size-based heterogeneities without the need for OMV fraction separation.

---

Membrane-bound nanostructures are ubiquitous across all domains of life. Extracellular vesicles (EVs), such as outer membrane vesicles (OMVs) produced by Gram-negative bacteria, are a notable example of these structures. OMVs are spherical, bound by a lipid bilayer, and composed of various components, including lipids, membrane proteins, lipopolysaccharides (LPS), and other biomolecules derived from the outer membrane of the bacteria^1–3^. OMVs transport a range of diverse cargo such as nucleic acids, proteins, and toxins, which play crucial roles in various biological functions of OMVs. While non-pathogenic bacterial OMVs share functions with eukaryotic EVs, such as cellular communication and removal of unwanted components, pathogenic bacterial OMVs have a unique role in transporting toxins to the host cell and facilitating the spread of disease^4–8^.

Although OMVs play a significant role in virulence and pathogenesis, certain features provide advantages for their use in various biotechnologies. Previously, an OMV based vaccine has been approved for use for meningococcal disease^9^, and their small size, biological nature, and diverse surface antigens make them attractive targets for the development of other vaccines as well^10,11^. Furthermore, recent studies have shown that OMVs can be bioengineered to deliver cytotoxic payloads directly to cancer cells, making them a promising tool for targeted cancer therapy^12,13^. Moreover, *Kim et al*. demonstrated that bioengineered OMVs can effectively reduce tumors even in the absence of cytotoxic payload, indicating their potential as a standalone therapeutic agent, further highlighting the potential of OMVs^14^. In addition OMVs have also shown promise as carriers for antibiotics, enzymes, and use as diagnostic agents^15–17^. Given the versatility and potential of OMVs, further research in this area is warranted to fully explore their potential in various fields.

Interestingly, OMVs originating from a single bacterial population can exhibit variations in size, protein composition, and encapsulated cargo^18,19^ These differences can be attributed to variances in the parent bacterium during growth. The intriguing heterogeneities observed in OMV structure and composition suggest that there may be significant differences in function and immunological properties between different types of OMVs. While the entry of OMVs into cells through various mechanisms has been widely acknowledged^20–23^, it was not until a recent study by *Turner et al*. that the influence of the size of *Helicobacter pylori* OMVs on their entry into epithelial cells was revealed^24^. Specifically, smaller OMVs (20-100 nm) enter through caveolin-mediated endocytosis, whereas larger OMVs (90-450 nm) enter through macropinocytosis and endocytosis^24^. Additionally, protein heterogeneity was noted within the OMV population, with smaller OMVs containing proteins that were absent in the larger ones^24^. These findings underscore the significance of investigating and comprehending the heterogeneity present within a population of OMVs.

Despite the potential significance of OMV heterogeneity in determining their function and potential applications in vaccine development and other biomedical areas, the current lack of research focused on characterizing the heterogeneities within an OMV population is a major limitation. This lack of research is primarily due to the limited analytical methods currently available for characterizing the heterogeneities within an OMV population. Ensemble assays such as ELISA and Western blots can provide information about the overall composition of OMV populations, however, they are unable to detect individual OMV heterogeneities^19,25–28^. In addition, to detect size-based heterogeneities, the OMV populations need to be separated using either the traditional density gradient ultracentrifugation^24^ or size exclusion chromatography (SEC) methods^29^ which can be time-consuming and add to the cost of the method. Recently, mass spectrometry has also been employed for OMV proteomics analysis^30–33^. However, it is important to note that this technique analyzes OMV ensembles, can be time and cost intensive, and it may necessitate specialized instrumentation. To fully comprehend OMV heterogeneity, it is necessary to use single-particle analysis methods, such as flow cytometry^24,34,35^ and electron microscopy^36–38^. However, these methods do come with certain limitations that can hinder their effectiveness. For example, flow cytometry operates on a continuous flow of OMVs, which makes it difficult to backtrack or perform additional analysis once heterogeneity is observed. Furthermore, flow cytometry requires special instrumental modifications to analyze nanoscale particles. Electron microscopy, on the other hand, can exhibit sample preparation artifacts and may not accurately represent the native structure of OMVs. To overcome the challenges posed by specialized equipment, high costs, and technical expertise required by existing OMV analysis techniques, there is an urgent need to develop new methods that can utilize general laboratory equipment. Investigating OMVs has the prospect to provide significant insights into bacterial pathogenesis, the development of new antimicrobial strategies, and the design of novel drug delivery systems.

Optical microscopy is a powerful tool to analyze bacterial outer membrane vesicles (OMVs)^20,22,28,39,40^ but it has a limitation where OMVs smaller than ∼200-400 nm appear as diffraction-limited spots and cannot be optically sized, due to which, any size-based heterogeneities can be masked. Previously, single particle fluorescence sizing has emerged as an alternative to determine the size of particles smaller than 200 nm^41–44^. In this letter, we present a platform for multi-parametric analysis of single OMVs using a general-purpose fluorescence microscope. Our method, illustrated in **(Figure 1)**, enables simultaneous detection of both the size and surface toxin/protein content on individual OMVs, providing a comprehensive characterization of these complex nanovesicles. Our approach involves a two-step process. First, we captured biotinylated OMVs on a streptavidin-passivated glass surface and characterized their size based on fluorescence intensity **(Figure 1a)**. Second, we used a toxin-specific antibody to categorize the OMVs into two groups: toxin-positive and toxin-negative and analyzed the size distribution of each group **(Figure 1b)**. This method is innovative in its ability to analyze single OMVs without the requirement of OMV fraction separation or specialized equipment, demonstrating its versatility and effectiveness.

**Figure 1:**
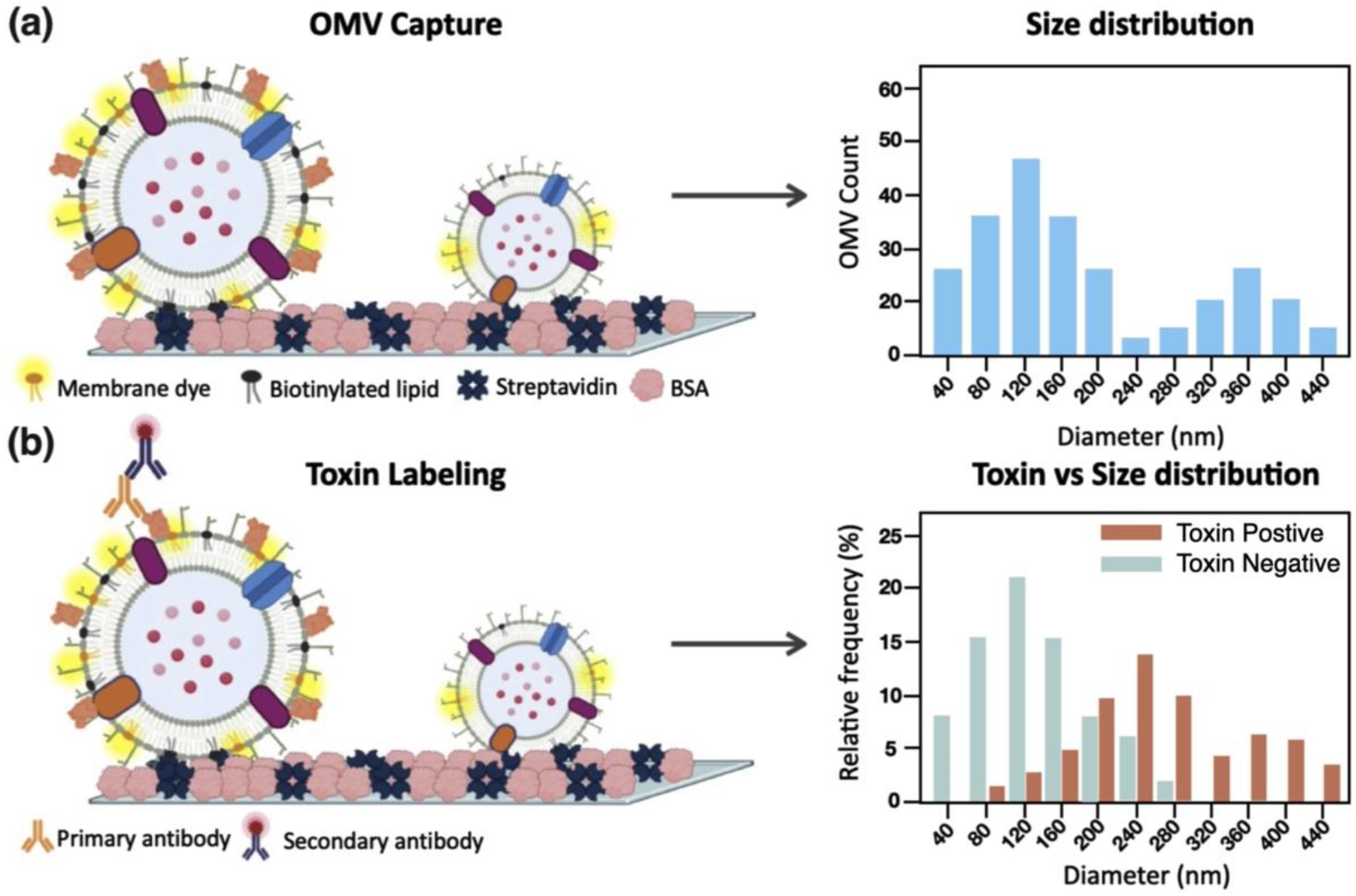
Schematic representation of the assay: **(a)** OMVs are biotinylated and fluorescently labeled with a membrane dye, immobilized on a glass surface, and their integrated fluorescence intensities are measured to generate their size distribution. **(b)** A toxin-specific antibody is introduced into the chamber and labeled with a fluorescent secondary antibody. The OMVs are categorized as toxin-positive and toxin-negative, and the size-based toxin distribution is determined.

To demonstrate the ability of our method to detect size-based toxin sorting, we utilized *Aggregatibacter actinomycetemcomitans (A*.*a*.), an oral bacterium associated with aggressive forms of periodontitis^45^. As part of its virulence, *A*.*a*. produces leukotoxin A (LtxA), which targets leukocytes^33,46^. *A*.*a*. secretes LtxA in two forms: water-soluble free LtxA and LtxA attached on the surface of OMVs^36^. Interestingly, *A*.*a*. produces a bimodal size distribution of OMVs, where a predominant population of ∼100 nm in diameter is observed along with a minor population of ∼350 nm in diameter^36^. Here, we aimed to determine how the toxin was distributed amongst the OMV populations.

First, we aimed to analyze the diameter of OMVs produced by two *A*.*a*. strains, JP2 and AA1704. While JP2 secretes high levels of leukotoxin, AA1704 is an isogenic mutant of JP2 that is deficient in LtxA production^47–49^. Dynamic light scattering (DLS) analysis reveals that both strains produce a heterogenous population of OMVs, as indicated by the bimodal size distribution observed **(Figure 2a)**. Our next objective was to determine the size of OMVs using single-particle fluorescence sizing. Labeling OMVs with fluorescent lumen dyes is one possible approach, but it presents challenges such as incomplete labeling of OMVs and uneven packing of the dye. To obtain an accurate representation of size using fluorescence microscopy, we utilized a lipophilic membrane dye (DiI) that incorporates itself into the OMV membrane, providing a reliable representation of the surface area. After exposing OMVs to Biotin Cap-PE, which incorporates into the membrane due to its hydrophobic nature, we captured them on streptavidin- and BSA-passivated glass **(Figure 2b)**. We opted to employ the biotin-streptavidin immobilization approach, as it offers a high affinity between biotin-streptavidin, guaranteeing firm immobilization of OMVs throughout the sizing and heterogeneity experiments. Furthermore, this method enables capture without any size or surface marker bias. We preceded by measuring the fluorescence intensity of the captured OMVs and discovered that each OMV displayed a unique integrated fluorescence intensity, which is proportional to its surface area, and thus can be used to determine the size distribution of the vesicle population **(Figure S1)**. Building upon the approach pioneered by Stamou and coworkers^41–44^, the intensity was then transformed into size information, creating a fluorescence OMV size distribution **(Figure 2c)**. For more information on the size determination process, please refer to the SI methods section titled ‘Size determination using fluorescence integrated intensities’. The analysis revealed the presence of two populations of OMVs, one with a diameter of roughly 100 nm and the other with a diameter of around 300 nm. The results obtained from the fluorescence intensity analysis were consistent with our previous findings from DLS.

**Figure 2:**
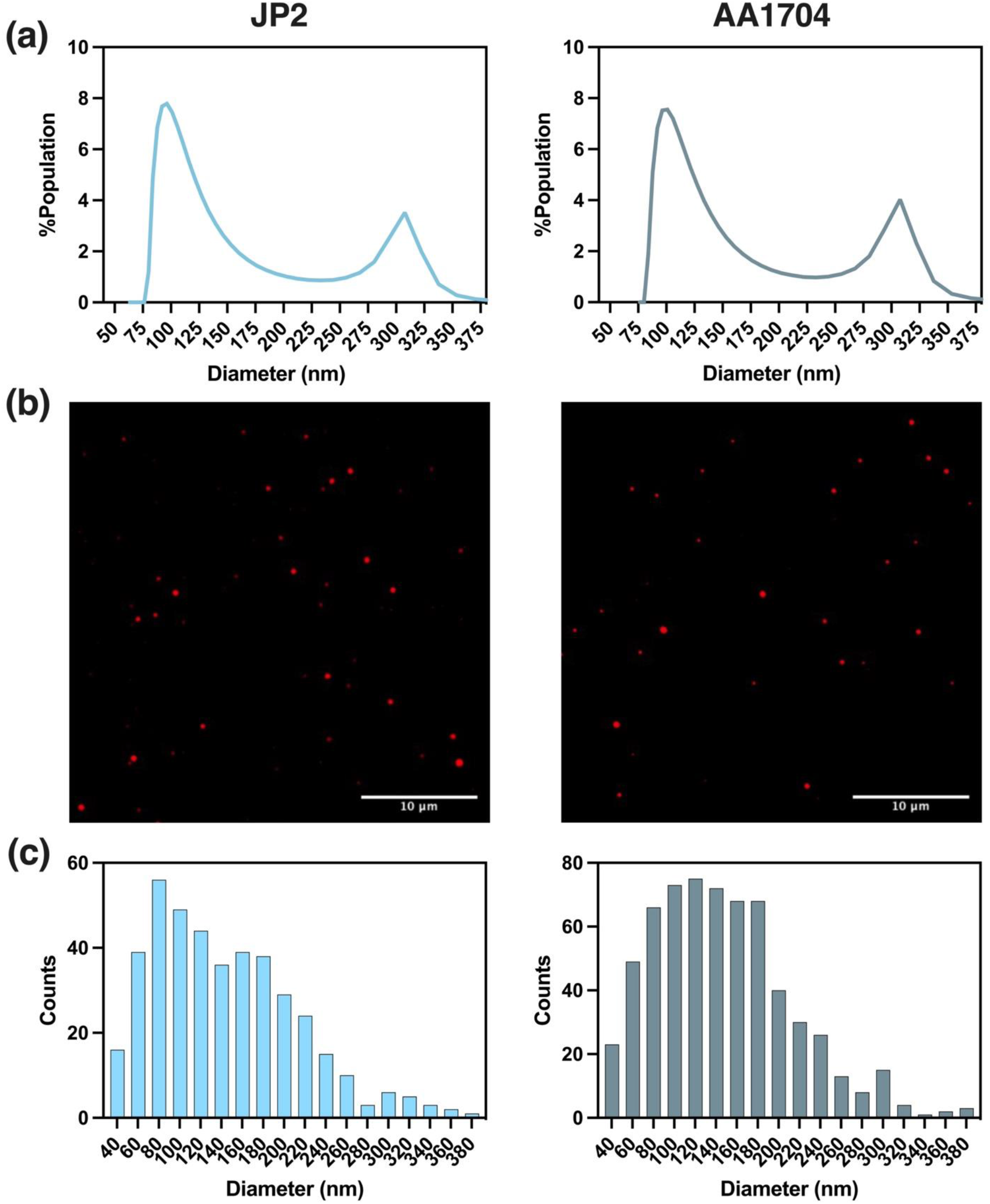
*A*.*a*. OMV characterization: **(a)** bimodal size distribution by DLS analysis. **(b)** Integrated fluorescence intensity of DiI-labeled individual OMVs **(c)** Bimodal size distribution by fluorescence particle sizing analysis. A total of 646 individual OMVs were counted for AA1704 and 416 OMVs were counted for JP2.

Previously the fluorescence single particle sizing has been utilized for synthetic particles. Therefore, there is a possibility that the bimodal size observed in *A*.*a*. OMVs may not be an accurate representation of their size, but rather a result of biological factors. To confirm that the bimodal size distribution of *A*.*a*. OMVs truly reflected their size heterogeneity, we also analyzed liposomes and *E. coli* OMVs, both of which have a unimodal size distribution **(Fig. S2)**. Both *E. coli* OMVs and *A*.*a*. OMVs were labeled with the same membrane fluorescent dye (DiI) and biotinylated to ensure a fair comparison of their integrated intensity. The results showed that the liposomes and *E. coli* OMVs had a unimodal integrated intensity distribution which results in a unimodal size distribution, confirming that the bimodal distribution observed in *A*.*a*. OMVs was due to their heterogeneous size population **(Figure S2)**. The integrated intensity distribution of *E. coli* OMVs displayed a distinct shift to lower values, indicating the presence of OMVs that are smaller than those produced by *A*.*a*. The majority of these OMVs had a square root integrated intensity of approximately 300 i.u. This observation suggests a correlation between the diameter of the OMVs and their intensity, as the smaller size of *E. coli* OMVs likely results in lower overall intensity. In contrast, *A*.*a*. OMVs displayed maxima at higher integrated intensities of 600 and 2250 i.u., suggesting a larger diameter for these OMVs **(Figure S2)**. These results highlight the effectiveness of fluorescence microscopy in accurately determining the size distribution of both heterogeneous and homogeneous size populations of OMVs.

Once the size of the OMVs was determined, we investigated the potential for toxin variation on the surface of *A*.*a*. OMVs. To examine this possibility, we conducted an experiment with a glass surface passivated by an anti-LtxA antibody **(Figure S3)**. The OMVs were then exposed to the passivated glass and counted based on the number of OMVs that bound in three 2048 × 2048 pixel areas. A control experiment was performed targeting outer membrane protein A (OmpA), which is commonly found on OMV surfaces, to normalize the count of OMVs **(Figure S3)**. Our results indicated a significantly fewer OMVs bound to the anti-LtxA passivation, suggesting that LtxA is present on only a subset of the OMVs **(Figure S3)**. Additionally, a control experiment was performed with the LtxA deletion mutant (AA1704), and no significant binding to the anti-LtxA was observed, further verifying the specificity of the antibody towards the toxin **(Figure S3)**.

To further investigate the observed discrepancy in the presence of the toxin on certain OMVs, we biotinylated the OMVs and immobilized them on a streptavidin surface. We then exposed the surface to the anti-LtxA antibody and added a secondary antibody conjugated with a fluorophore to track the presence of the toxin. We combined the channel displaying OMV membrane fluorescence and the antibody channel to visualize the colocalization between the two. This approach allowed us to determine the distribution of LtxA on the OMV surface and determine any size-based heterogeneities. Our results showed that while some JP2 OMVs were positive for the toxin, others lacked it **(Figure 3a)**. As a control, we tested for binding to outer membrane protein A (OmpA), which is present ubiquitously on the OMVs surface^28^, and observed non-discriminatory binding **(Figure 3b)**. As an additional negative control, we tested AA1704 OMVs for LtxA and did not observe any antibody binding **(Figure 3c)**. Our results confirm that that our platform is capable of detecting individual OMV heterogeneity, including size-based heterogeneities as well as compositional heterogeneities such as the presence of the toxin or proteins as demonstrated by the binding observed only in OMVs containing the toxin.

**Figure 3:**
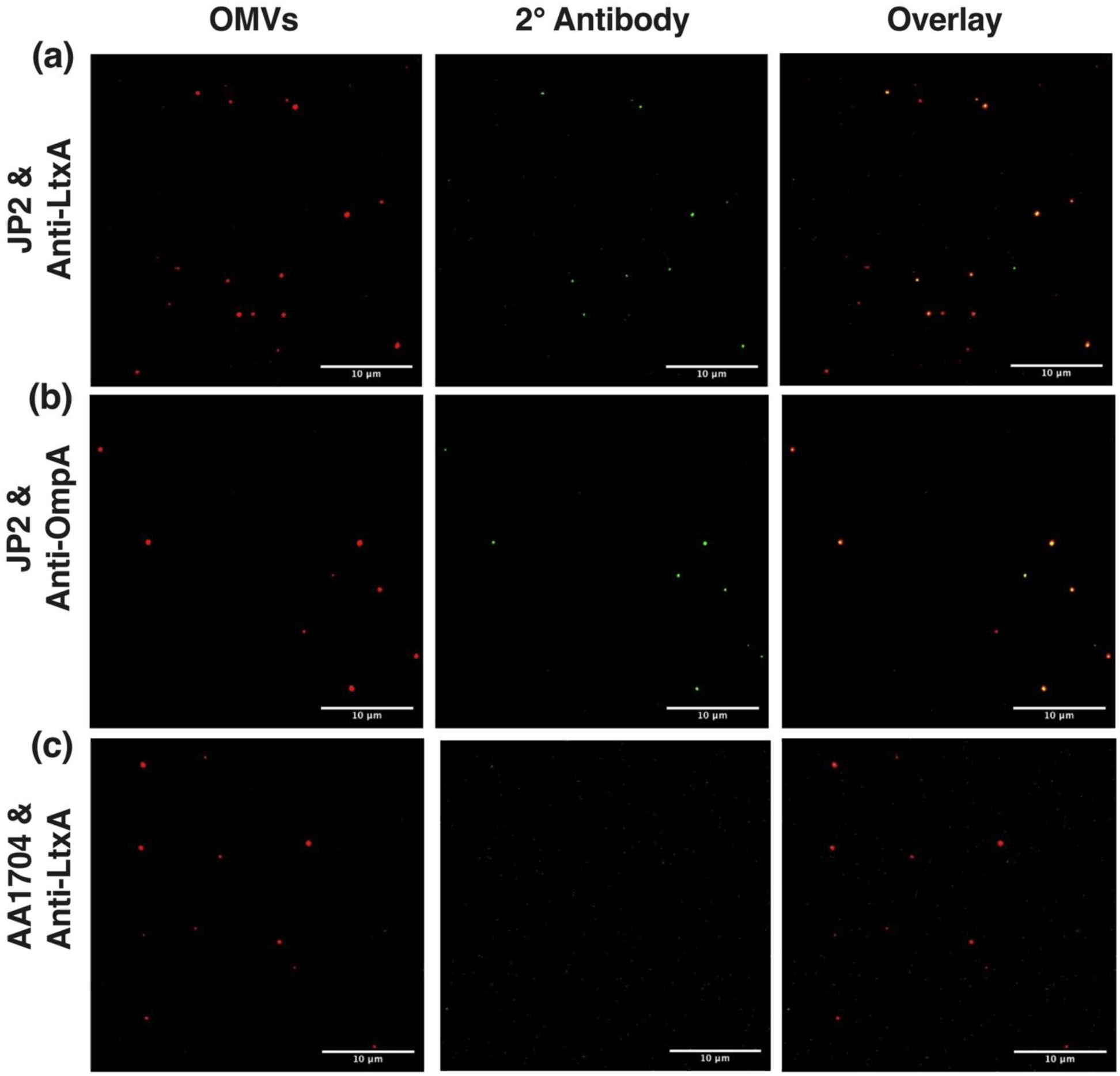
Surface protein analysis of *A*.*a*. OMVs: **(a)** OMVs from the JP2 strain were analyzed for the presence of LtxA. The fluorescence micrograph shows only a few OMVs with positive staining for LtxA, indicating that not all OMVs carry this virulence factor. **(b)** Positive control: OMVs were analyzed for ubiquitous OmpA. **(c)** Negative control: OMVs from the AA1704 strain, which is a leukotoxin deficient strain, were analyzed for the presence of LtxA.

To determine if the discrepancies in LtxA antibody binding with JP2 OMVs were size-based, we categorized the OMVs into two groups based on the presence of LtxA: toxin-positive and toxin-negative, and then analyzed their size distributions. Our findings indicate that LtxA was primarily present in larger OMVs, with smaller diameter OMVs devoid of the toxin. The significance of single OMV analysis was highlighted by our noteworthy findings: no LtxA-negative OMVs were found to have a diameter greater than 220 nm, and no LtxA-positive OMVs were found to have a diameter smaller than 60 nm **(Figure 4a)**. This analysis enables the detection of nanoscale discrepancies in OMVs that are often obscured by traditional ensemble assays. Furthermore, we investigated the correlation between toxin presence or absence and OMV diameter, and our observations revealed in the range of 140 nm, there was an even split of LtxA-positive and -negative OMVs, indicating a heterogeneous population. Our examination of overall OMV positivity and negativity for toxins in relation to diameter revealed a proportional increase in toxin-positive OMVs and a decrease in toxin-negative OMVs as OMV size increased **(Figure 4b)**.

**Figure 4:**
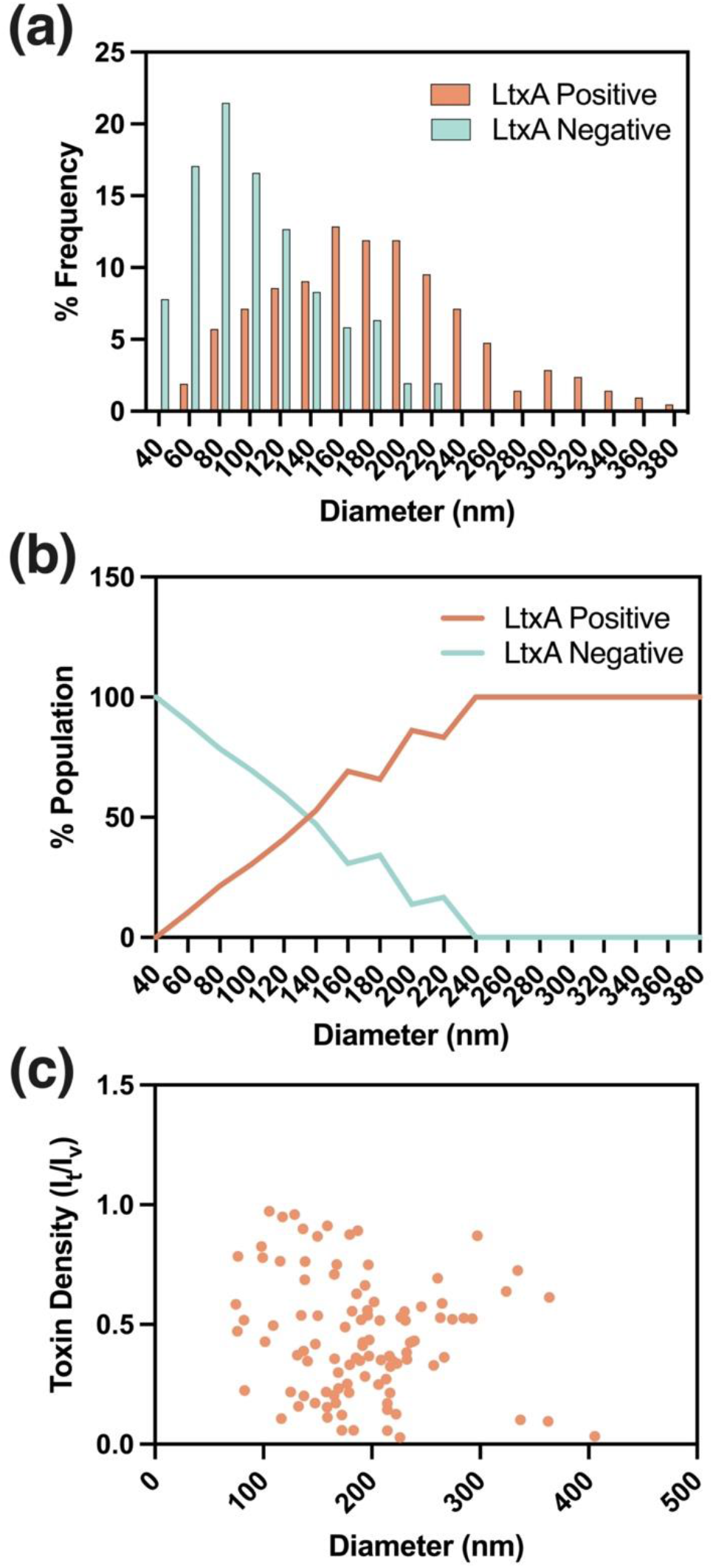
Size-based toxin heterogeneity analysis: **(a**) Fluorescently labeled OMVs were analyzed for the presence of LtxA toxin. No LtxA-negative OMVs were found to have a diameter greater than 220 nm, while no LtxA-positive OMVs had a diameter smaller than 60 nm.(**b)** Fraction LtxA positive and negative as a function of OMV diameter. At 140 nm, the proportion of OMVs containing the toxin and those being negative were found to be approximately equal. **(c)** Toxin density plotted against vesicle diameter. The Pearson correlation coefficient (r) of -0.168 and p-value of 0.906 indicate no relationship between the density of toxin and OMV size.

After observing the presence of LtxA in larger OMVs, we aimed to determine if there was a correlation between the size of OMVs and the density of the toxin. To do this, we analyzed the toxin density where we divided the intensity of the LtxA antibody by the intensity of the vesicle membrane **(Figure 4c)**. We found that the size of OMVs and the density of the toxin were not significantly correlated, as supported by a Pearson correlation coefficient (r) of -0.168 and a p-value of 0.906. This suggests that the larger OMVs, which have 10 times more surface area than the smaller OMVs, do not necessarily have more LtxA present per vesicle. These results highlight the importance of further investigation into the mechanisms controlling toxin sorting and distribution within OMVs.

We validated our toxin discrepancy analysis of OMVs using an ELISA and western blot assay. To investigate size-based heterogeneity, it was crucial to separate the OMVs based on size prior to the western blot analysis. An ultra-fine Sepharose S1000 column has been used previously for size-based OMV separation^26^, which is a simpler alternative to traditional methods like density gradient centrifugation. We aimed to utilize this method to separate *A*.*a*. OMV subpopulations based on their size. Our analysis of chromatographic fractions confirmed the presence of two populations of OMVs, eluting in different volumes (**Figure 5a**). DLS measurements showed that larger OMVs (over 250 nm) were found in the 34-36 mL fractions, while smaller OMVs (under 250 nm) were found in the 44-52 mL fractions **(Figure 5b)**.

**Figure 5:**
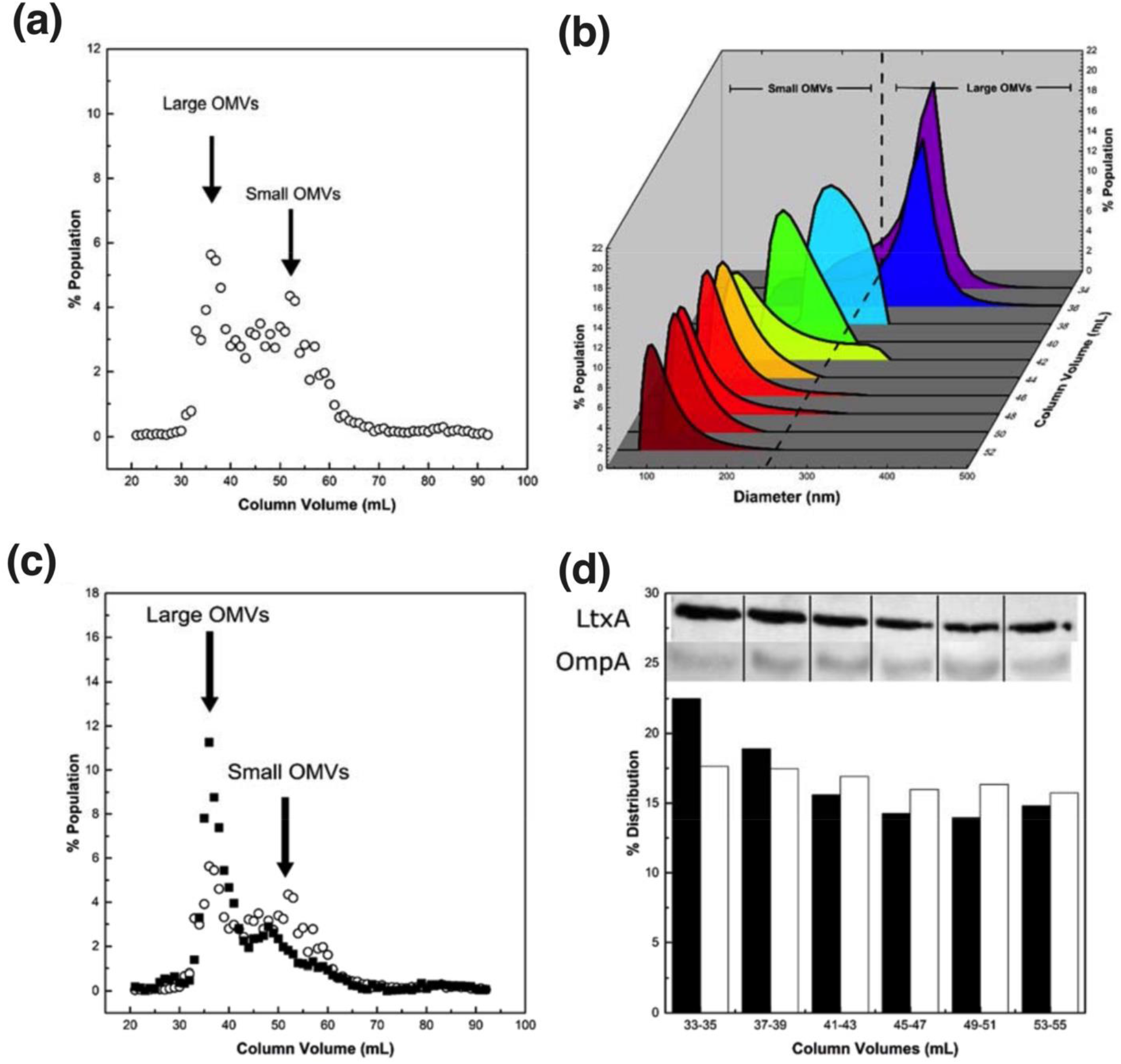
Analysis of OMVs separated based on their size: **(a)** OMVs were separated by SEC. The fluorescence of a lipid probe, FM 4-64 (open circles), was used to identify fractions containing OMVs. **(b)** SEC fractions were analyzed using DLS to determine the size distribution of OMVs. **(c)** The concentration of LtxA toxin in each fraction is measured using ELISA (black squares) while the lipid concentration in each fraction is determined using FM 4-64 dye (open circles). **(d)** Western blot analysis of LtxA and OmpA in each fraction, normalized to their respective lipid content.

Next, the OMV fractions were analyzed to determine the presence of LtxA. The results from ELISA showed that much of the LtxA was found in fractions containing the larger OMVs, (**Figure 5c**), in agreement with the single-particle fluorescence analysis. To address the potential discrepancy of the toxin being observed in larger OMVs due to a greater surface area, we performed ELISA on SEC fractions that were normalized by their total lipid concentration. The results showed that the larger OMVs (33-35 mL) had more LtxA, and the smaller fractions had less **(Figure 5c)**. As a positive control, the presence of OmpA in the OMV fractions was also detected by western blotting, and the results showed that OmpA was present in all fractions, while a notable trend of increasing LtxA level with OMV size was observed (**Figure 5d**). The ELISA and western blot results served to confirm the validity of our individual OMV fluorescence approach.

OMVs are considered an essential component in many biological systems and play a vital role in various processes, including cell-to-cell communication, horizontal gene transfer, and pathogenesis. Despite their significance, the heterogeneous size and composition of OMVs pose challenges in understanding their functionality and interaction with target cells. Conventional OMV analysis assays, which rely on population measurements, ignore variations between individual OMVs. Although single OMV analysis methods exist, they can often be limited in their scope.

Our study presents a novel solution using fluorescence particle sizing to size and uncover size-based heterogeneities in bacterial OMVs. This method enables simultaneous detection of size and surface toxin/protein content on individual OMVs, providing a comprehensive characterization of these complex nanoparticles. Our results demonstrate the versatility and effectiveness of our method in accurately determining the size distribution and heterogeneity of OMVs. We utilized fluorescence microscopy to analyze heterogeneous populations of OMVs produced by the oral bacterium *A*.*a*. The results from our study provide new insights into the size distribution of *A*.*a*. OMVs and reveal the presence of two populations of OMVs, one with a size of roughly 100 nm and the other with a diameter of around 300 nm. Furthermore, our single particle analysis allowed us to discover that no LtxA-negative OMVs were found to have a diameter greater than 220 nm, and no LtxA-positive OMVs were found to have a diameter smaller than 60 nm. These results highlight the significance of single OMV analysis as it enables the detection of nanoscale discrepancies in OMVs that are masked by traditional ensemble assays.

Previous studies have mainly reported unimodal size distributions for clinically significant OMVs, including those released by *Acinetobacter baumannii, Pseudomonas. aeruginosa, Escherichia coli, and Enterobacter cloacae*^51–54^. It is important to note that although most OMVs exhibit a unimodal size distribution, they still have a wide size range. Our results highlight that the first size distribution of *A*.*a*. OMVs, ranging from 75 – 175 nm, contained a mixture of toxin-positive and toxin-negative vesicles suggesting that there can be substantial differences within a unimodal size distribution OMV, leading to function differences as previously seen with the bimodal OMVs of *H. pylori*^24^.

In conclusion, our platform is unique in its approach as it does not require OMVs to be separated and is non-invasive, utilizing a general-purpose microscope instead of specialized equipment. Our study highlights the versatility and effectiveness of our method in accurately determining the size distribution and heterogeneity of OMVs and offers a more comprehensive and advanced approach to studying OMVs.

## Supporting information

supplemental meterials

## Acknowledgments

This work was supported by grants from the National Institutes of Health to N.J.W. (R21GM134414) and A.C.B. (R21DE025275). N.J.W. and A.C.B. also acknowledge support from a Lehigh University CORE Grant. Illustrations were created using Biorender.com.

