## supplemental meterials for "Identifying size-dependent toxin sorting in bacterial outer membrane vesicles"

**Running title: Size dependent toxin sorting**

**Keywords: Single particle sizing, fluorescent microscopy, outer membrane vesicles, toxins**

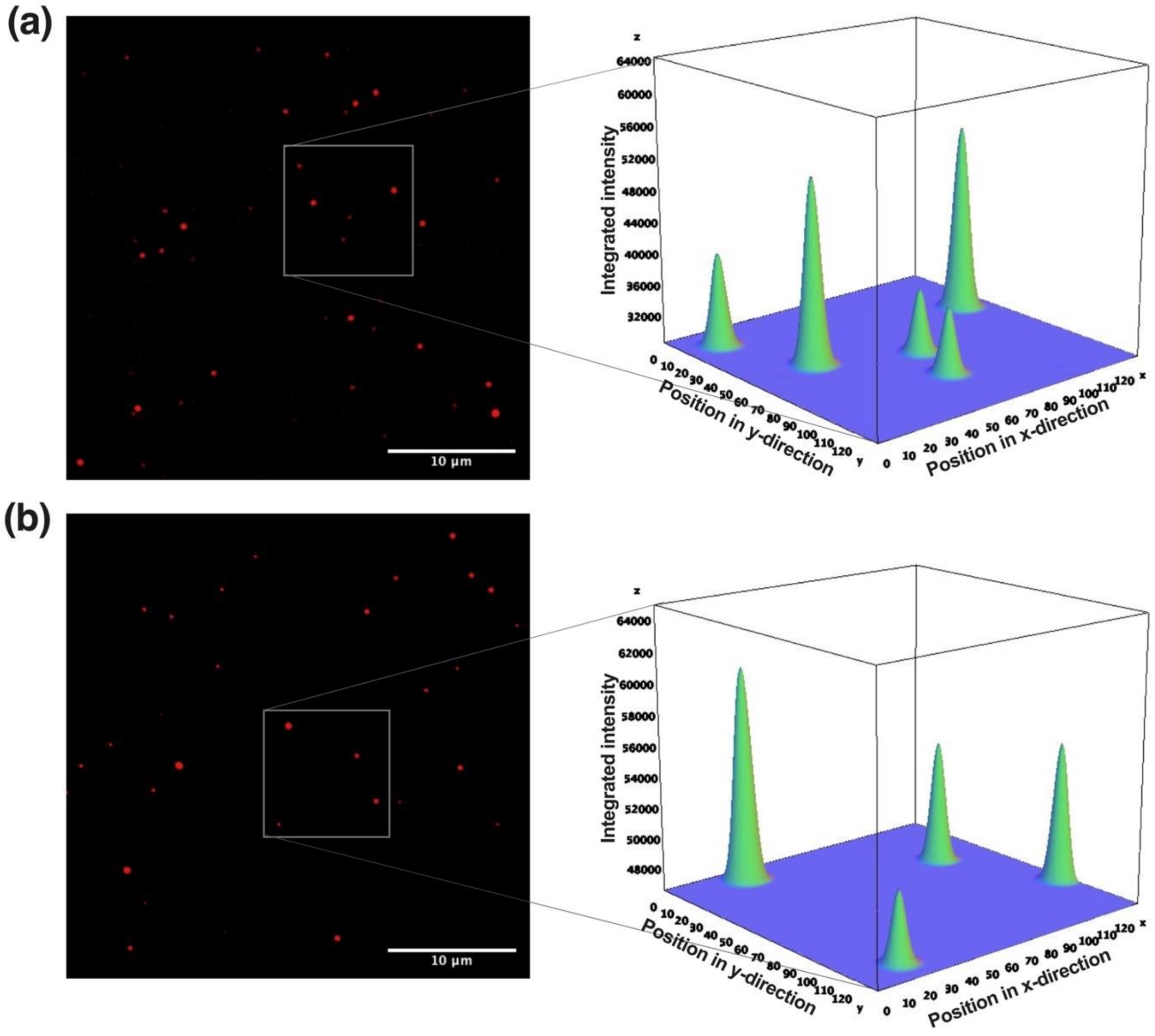


**Figure S1: Quantifying the integrated fluorescence intensity of immobilized OMVs:** Fluorescence micrographs of OMVs stained with a membrane label (DiI) and intensity profiles for OMVs from **(a)** JP2 and **(b)** AA1704 strains.

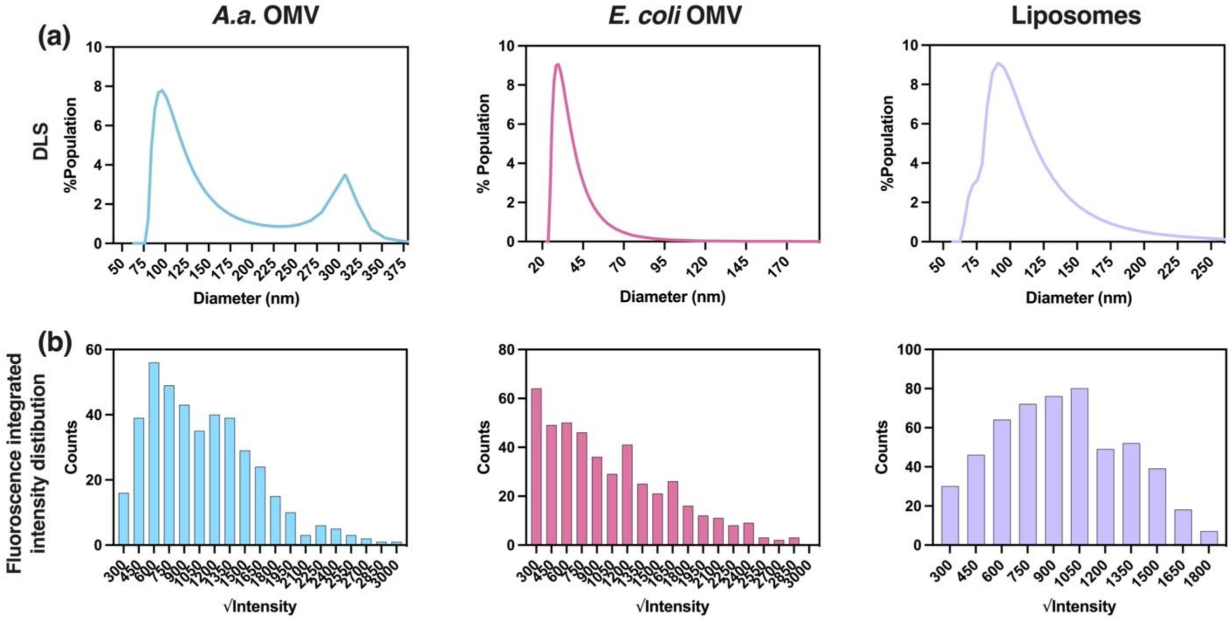


**Figure S2: comparative analysis of size and fluorescence intensity for *A.a.* OMVs, *E. coli* OMVs, and liposomes**. **(a)** DLS size distribution for each type of particle. **(b)** Fluorescent integrated intensity distribution for each type of particle. Integrated intensity distributions are proportional to the distribution of radii.

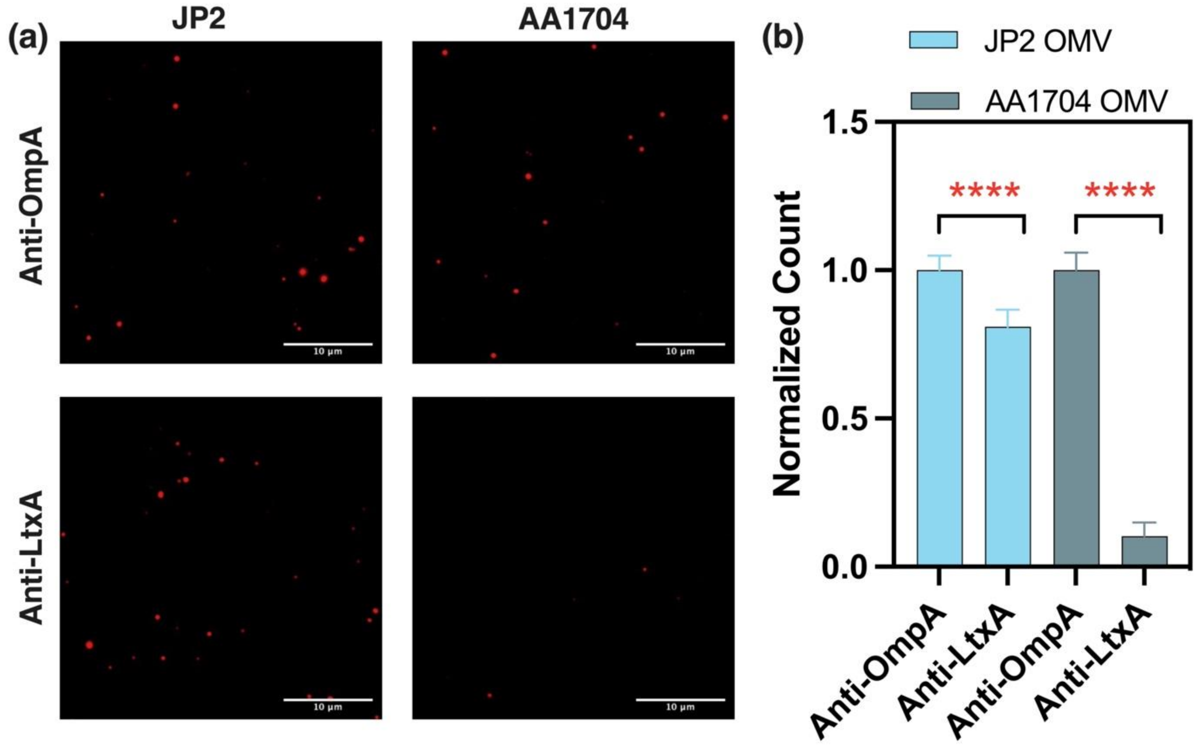


**Figure S3: Analysis of OMV surface proteins: (a)** Micrographs of OMVs captured on glass passivated with anti-LtxA antibody, compared to ubiquitous anti-OmpA protein passivation. **(b)** The normalized counts of OMVs show significantly lower binding of JP2 OMVs to the anti-LtxA antibody compared to the negative control, AA1704 OMVs. ****A p-value of less than 0.0001 indicates a significant difference in the presence of LtxA on the OMV surface.

**Methods:**

***Reagent and chemicals:***

Following lipids used in this study were bought from Avanti polar lipids: 1-palmitoyl-2-oleoyl-glycero-3-phosphocholine (POPC), GM1 (ovine brain), 1,2-dioleoyl-*sn*-glycero-3-phosphoethanolamine-N-(Biotin Cap-PE), and 1,2-dioleoyl-*sn*-glycero-3-phosphoethanolamine-N-(Cyanine 5). Streptavidin was purchased from New England Biolabs. Fatty-acid-free Bovine serum albumin (BSA) and NeutrAvidin was purchased from Sigma-Aldrich. Anti-LtxA antibody was collected from the supernatant of hybridoma cell line LTA83[5], gifted by Dr. Edward T. Lally, University of Pennsylvania, grown in serum-free medium; the antibody was purified using a protein-G column. Anti-OmpA rabbit polyclonal antibody was purchased from Antibody Research Corporation. Secondary antibody with Alexa-488 label was purchased from Abcam. Dimethyl sulfoxide (DMSO) was purchased from Sigma-Aldrich. DiI and DiO membrane labels were purchased from Invitrogen, ThermoFisher. All experiments were performed in Tris buffer composed of 150 mM NaCl and 10 mM Tris-base (pH 7.0).

***Liposome preparation:***

Appropriate molar ratios of desired lipids were dissolved in chloroform, or a chloroform/methanol mixture; the solution was mixed in a glass vial and the solvent was evaporated to obtain thin lipid films. The lipid films were rehydrated in Tris I buffer with a final concentration of 1 mg/mL, followed by gentle vortexing and bath sonication for 10 minutes. The resulting liposomes were extruded through a 100 nm polycarbonate membrane (Cytiva Life Sciences) using a Mini-Extruder (Avanti polar lipids) with 23 passes.

***OMV membrane labeling and biotinylation:***

To label the OMVs with fluorescent membrane dye, we utilized the lipophilic membrane dye, DiI. 20 µM of DiI dissolved in ethanol was added to the OMVs and incubated for one hour at 37°C to ensure optimal labeling efficiency. Additionally, we biotinylated the OMVs using biotinylated Cap-PE. To achieve this, we prepared Biotinylated Cap-PE in DMSO and then added 20 µM of the stock to the OMVs, which were then incubated at 37°C for one hour.

***Purification of A.a. OMV:***

*A.a.* strains JP2 and AA1704 were grown in trypticase soy broth (30 g/L, BD Biosciences, Franklin Lakes NJ) and yeast extract (6 g/L, BD Biosciences), supplemented with 0.4% sodium bicarbonate (Fisher Scientific, Hampton, (NH,) USA), 0.8% dextrose (BD Biosciences), 5 µg/mL vancomycin (Sigma-Aldrich, St. Louis, (MO,) USA), and 75 µg/mL bacitracin (Sigma-Aldrich). A starter culture was grown for 16 hr in a candle jar, then inoculated into a larger culture and allowed to grow for 24 hr at 37°C. The bacteria were centrifuged at 10,000 × g for 10 min, then again at the same speed for 5 min, followed by filtration through a 0.45 µm filter. This supernatant was then ultracentrifuged at 105,000 × g for 30 min, resuspended in PBS (pH 7.4) and ultracentrifuged again. The final pellet was resuspended in PBS.

***Dynamic Light Scattering (DLS) of Liposomes and OMV:***

Liposome and OMV diameters were measured using dynamic light scattering (DLS). OMVs were diluted to 1:100 dilution factor and the liposomes were diluted to a final concentration of 0.25 mg/mL in Tris buffer. DLS measurements were collected using the ALV/CGS-3 Compact Goniometer System spectrometer. The data was collected in triplicate for 120 seconds at the wavelength of 623.8 nm and a 90° scattering angle. ALV software number-weighted regularized fit with an allowed membrane thickness (r*) of 5 nm was used to determine the size distribution. A weighted average was calculated to obtain the mean radius for fluorescence microscopy particle sizing.

***Liposome and OMV capture:***

Glass coverslips (cleaned with IPA and 2% SDS) were UV/Ozone treated for 10 min (UV/Ozone ProCleaner Plus, BioForce Nanosciences) and passivated with 200 µg/mL of NeutrAvidin to capture liposomes or streptavidin to capture OMVs, prepared in 1% BSA. The coverslip was washed with 1% BSA, and either 0.5 µg/mL of liposomes or OMVs (1:50 dilution) were added to the chamber and incubated for 1 hour. Unbound liposomes or OMVs were washed from the chamber, and subsequently imaged using an Nikon Ti inverted microscope equipped with a 100× oil immersion objective. LED light engine from Aura II, Lumencor was used to excite fluorescence and a Cy5 (Chroma) or a TRITC (Chroma) filter set was utilized. A 2048 × 2048 pixel sCMOS camera (Orca Flash 4.0 v2, Hamamatsu) was used to capture all images. Image J was used to analyze the images and calculate the integrated intensity of the OMVs. The fluorescence intensity was converted to a size using the equations described above.

***OMV heterogeneity analysis:***

Once the OMVs were imaged, a full-length anti-LtxA antibody[5] was introduced to the chamber (1:1000 dilution). The anti-LtxA antibody was incubated for 1 hour and subsequently washed with Tris buffer. 2 µg/mL of secondary IgG with Alexa-488 (Abcam) was added to the chamber and incubated for 1 hour. Excess antibody was washed, and the samples were imaged, as described previously. The FITC (Chroma) filter set was used to capture the antibody binding. Using Image J, the TRITC and FITC color channel were overlayed, and the x/y coordinate of the OMVs and antibody were analyzed to determine toxin-positive/negative OMVs. The toxin-positive and toxin-negative OMVs were separated into two categories, and their size distribution was plotted using GraphPad Prism v 9.0. To determine the toxin density present on the surface of the OMVs, the integrated intensity from the antibody was divided by the integrated intensity of the respective OMVs ($I_t/I_v$).

***Antibody passivation for OMV surface heterogeneity:***

Clean glass coverslip (2% SDS and UV-ozone treated) were passivated with either Anti-OmpA antibody or Anti-LtxA antibody (1:100 dilution) prepared in 1% BSA and incubated for 1 hour. The chamber was washed with 1% BSA and OMVs (1:50) dilution were added. The unbound OMVs were washed thoroughly with Tris buffer and subsequently imaged using an Nikon Ti inverted microscope, equipped with a 100× oil-based immersion objective. LED light engine from Aura II, Lumencor was used to excite fluorescence and a TRITC (chroma) filter set was utilized. A 2048 × 2048 pixel sCMOS camera (Orca Flash 4.0 v2, Hamamatsu) was used to capture all images. Using Image J, OMVs were counted over three 2048 × 2048 pixel frames (repeated 3 times) and data were normalized to the Anti-OmpA passivated OMV count. The data were analyzed using GraphPad Prism v 9.0.

***SEC of A.a. OMV***

SEC was used to separate OMVs by size; a 1.5 cm x 50 cm (bed volume 85 mL) was packed with Sephacryl™ S-1000 superfine resin (GE Healthcare, Chicago, IL, USA) and equilibrated with two bed volumes of PBS. A 2-mL OMV sample was loaded, eluted with PBS, and one-mL fractions were collected. Fractions were analyzed for lipid and LtxA content, as described below.

The percentage of lipid in each SEC fraction was measured using the FM 4-64™ dye (ThermoFisher Scientific, Waltham, MA). First, 50 μL of each fraction was incubated with FM 4-64™ (0.1 mg/mL) for 15 sec. Following incubation, the fluorescence of the sample was measured on a Tecan plate reader with an excitation wavelength of 515 nm and an emission wavelength of 640 nm. The fluorescence intensity of each fraction was divided by the summed intensities to calculate a percentage of total lipid in each fraction.

The LtxA concentration in each SEC fraction was measured using an ELISA. Fractions were incubated in a MaxiSorp Immuno 96-well plate (ThermoFisher Scientific) for 3 hr, washed five times with ELISA wash buffer (25 M Tris, 150 mM sodium chloride, 0.1% fatty-acid free bovine serum albumin (BSA)) then blocked in 1% BSA in the same buffer. The plate was then incubated with anti-LtxA antibody[5] in 1% BSA/buffer overnight at 4 °C. Following five washes with ELISA wash buffer, the plate was incubated in goat anti-mouse horseradish peroxidase (GAM-HRP) at a 1:5000 ratio (SouthernBiotech, Birmingham, AL). Lastly, the plate was imaged using 1-Step™ Ultra TMB ELISA substrate solution (ThermoFisher Scientific) until signal appeared, then the reaction was stopped using 2 M sulfuric acid. The absorbance at 450 nm was measured on a Tecan plate reader. The resulting absorbance of each fraction was divided by the summed absorbances to calculate a percentage of total LtxA in each fraction.

LtxA content was also analyzed by western blotting. Sodium dodecyl sulfate polyacrylamide gel electrophoresis (SDS-PAGE) was performed using 7.5% acrylamide gels. Western blotting for LtxA was accomplished by transferring the proteins to a nitrocellulose membrane overnight. The blots were washed three times in tris-buffered saline with 0.1% tween (TBST) and then blocked with blotto solution (5% dried milk in TBST) for 1 hr. LtxA was then detected using a monoclonal anti-LtxA antibody[5] overnight at 4°C, followed by GAM-HRP for 1 hr. The blot was imaged using SuperSignal™ West Dura substrate (ThermoFisher). To measure the amount of outer membrane protein-A (OmpA) in each fraction, a similar procedure was used, with the anti-Gram-negative OmpA antibody (111228, Antibody Research Corporation, St. Charles, MO) at a final concentration of 0.24 μg/mL, followed by goat anti-rabbit horseradish peroxidase (GAR-HRP, SouthernBiotech). Densitometry analysis on the blots was accomplished using ImageJ.

***Size determination using fluorescence integrated intensities:***

Optical microscopy is a useful tool for studying membrane-bound structures, but it has a limitation where particles smaller than ~200-400 nm appear as diffraction-limited spots and cannot be optically sized. To overcome this limitation, previous studies have utilized the integrated intensities of microscopic spots to determine the size of individual particles and liposomes[1–4] . To demonstrate that fluorescence microscopy can be used to determine the size of membrane bound structures, we aimed to size liposomes of known diameter. Biotinylated liposomes were immobilized on a streptavidin passivated glass surface, and their fluorescence intensities were analyzed **(Figure S4a)**. To enhance the threshold of our analysis, we used the

ImageJ particle analysis function and set a threshold value to differentiate the OMVs from the background. We then used the ImageJ particle analysis function to obtain the counted particles' integrated intensity **(Figure S4b)**. Individual liposomes exhibit different integrated intensities, which were used to determine their sizes **(Figure S4c)**. To convert the intensity distribution to a diameter distribution, the following mathematical conversions were used. The surface area of the liposomes ($SA_{Lip}$) is directly proportional to the integrated fluorescence intensity ($I_M$), meaning that larger OMVs will exhibit stronger fluorescence (Eq. 1.1). By assuming that liposomes are spherical, we were able to solve for their surface area (Eq 1.2). Eq. 1.1 and 1.2 can be combined to relate the intensity observed to the radius (r) using a correlation factor $k$ (Eq. 1.3). This correlation factor *k* relates the intensity of a single vesicle to its radius in nm, and it is dependent on factors such as the photophysical properties of the fluorophore and the signal detection efficiency. *k* can be calculated using (Eq. 1.4), where $r_{mean}$ is the mean radius determined using other sizing methods. In this study, DLS was used to calculate the $r_{mean}$, and the radius can be found using (Eq. 1.5), and a size distribution is generated.

$$SA_{Lip} \propto I_M \qquad \textbf{(Eq. 1.1)}$$

$$SA_{Lip} = 4\pi r^2 \qquad \textbf{(Eq. 1.2)}$$

$$r = k\sqrt{I_M} \qquad \textbf{(Eq. 1.3)}$$

$$k = \frac{r_{mean(DLS)}}{\sqrt{I_M}_{(Mean)}} \qquad \textbf{(Eq. 1.4)}$$

$$r = \frac{r_{mean(DLS)}}{\sqrt{I_M}_{(Mean)}} * \sqrt{I_M} \qquad \textbf{(Eq. 1.5)}$$

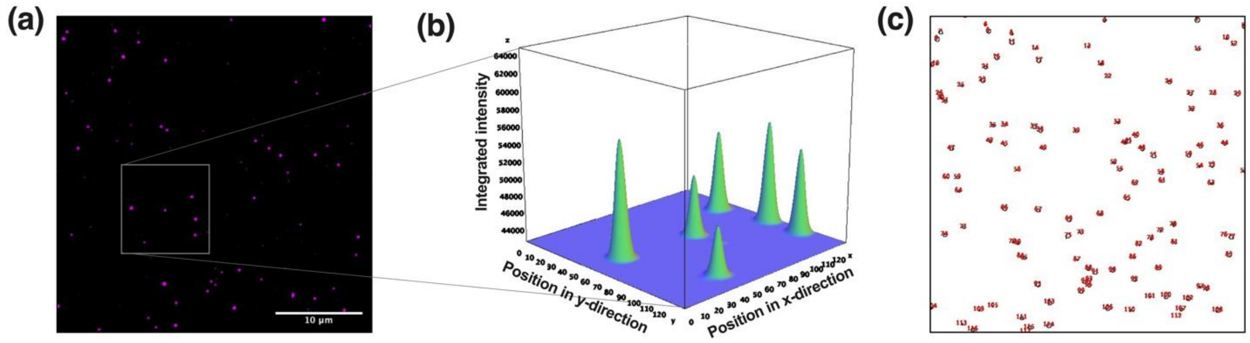


**Figure S4: (a)** Biotinylated liposomes were immobilized on a streptavidin passivated glass surface. **(b)** Liposomes were identified using ImageJ particle tracking analysis and their raw integrated intensity was obtained. **(c)** A 3-D intensity profile plot was generated to demonstrate that each liposome contains a unique integrated intensity despite being in proximity to each other.
